# The mitochondrial genome of the hammerhead flatworm *Bipalium nobile* and its phylogenetic implications

**DOI:** 10.64898/2026.05.31.729009

**Authors:** Masahiro Omura, Soma Tomihara, Ryuhei Minei, Daiki Haraguchi, Shuichi Wada

## Abstract

We sequenced the nearly complete mitochondrial genome of the hammerhead flatworm *Bipalium nobile* Kawakatsu and Makino, 1982 using short-read sequencing technology, yielding a 16,018 bp genome comprising 12 protein-coding genes, 22 tRNA genes, and 2 rRNA genes. The composition and order of genes were consistent with those observed in the closely related species *Bipalium kewense* and *Diversibipalium multilineatum*, except for the position of tRNA-Glu. Phylogenetic analysis based on all mitochondrial proteins from species within the family Geoplanidae supports the monophyly of a clade comprising *B. nobile, B. kewense*, and *D. multilineatum*. The mitochondrial genome sequence obtained in this study provides a valuable resource for investigating the genetic diversity and population structure of *B. nobile*, a soil-dwelling predator with the potential for global spread as an invasive organism.

## 1. Introduction

Hammerhead flatworms (Platyhelminthes, Geoplanidae, Bipaliinae) are soil-dwelling animals characterized by an elongate body and a distinctive semilunar head. As predators of land snails and earthworms, they occupy a high trophic position in soil ecosystems. In recent years, several species have expanded their ranges as non-native organisms, raising concerns about their ecological impacts. However, these impacts remain poorly understood, highlighting the need for further studies on the taxonomy and genetic diversity of hammerhead flatworms. Mitochondrial genome sequencing provides a useful resource for such taxonomic and population genetic analyses. To date, mitochondrial genomes of seven species of the subfamily Bipaliinae have been deposited in public databases.

Among hammerhead flatworms, three closely related species with longitudinal stripes and relatively large body sizes deserve particular attention: *Bipalium nobile, B. kewense*, and *Diversibipalium multilineatum. D. multilineatum* is placed in the collective group *Diversibipalium*, a genus established for species whose reproductive organs and other diagnostic characters have not been examined. Molecular phylogenetic analyses indicate that it forms a monophyletic clade with *B. kewense* and *B. nobile*, supporting a close relationship among the three species.

*B. kewense*, native to Southeast Asia, has spread to more than 78 countries worldwide (Gastineau et al. 2025). *D. multilineatum*, native to Japan, has been recorded from 10 European countries (de Waart et al. 2025). In contrast, *B. nobile* is thought to originate from China and has been reported only from Japan and South Korea. Despite their close phylogenetic relationships, these species differ markedly in the extent of their range expansion beyond their native distributions. Comparative studies of their biology and biogeographic history may provide insight into the mechanisms underlying range expansion and invasion success in hammerhead flatworms.

The mitochondrial genomes of *B. kewense* and *D. multilineatum* have been reported, but that of *B. nobile* has not. In this study, we determined a near-complete mitochondrial genome sequence of *B. nobile* from an individual collected in Japan. Further analyses based on mitochondrial genome data from all three species will help clarify their population structure and provide insight into the history of their range expansion.

## 2. Materials and methods

*B. nobile* was captured at the base of Mt. Tamura, Nagahama City, Shiga Prefecture, Japan (latitude: 35°21′42″ N, longitude: 136°17′24″ E) on 13 June 2024 (Figure 1). Genomic DNA was extracted from the dissected caudal portion using the Puregene Tissue Core Kit (QIAGEN, Hilden, Germany). The sequencing library was constructed using MGIEasy FS DNA Library Prep Set (MGI, Shenzhen, China) and sequenced with the DNBSEQ-G400RS (2 × 150 bp) at Genome-Read (Kagawa, Japan). The remaining part of the specimen was deposited at the Lake Biwa Museum (website: https://www.biwahaku.jp/; contact person: Takahito Suzuki) under the voucher number LBM1340000206.

**Figure 1.**
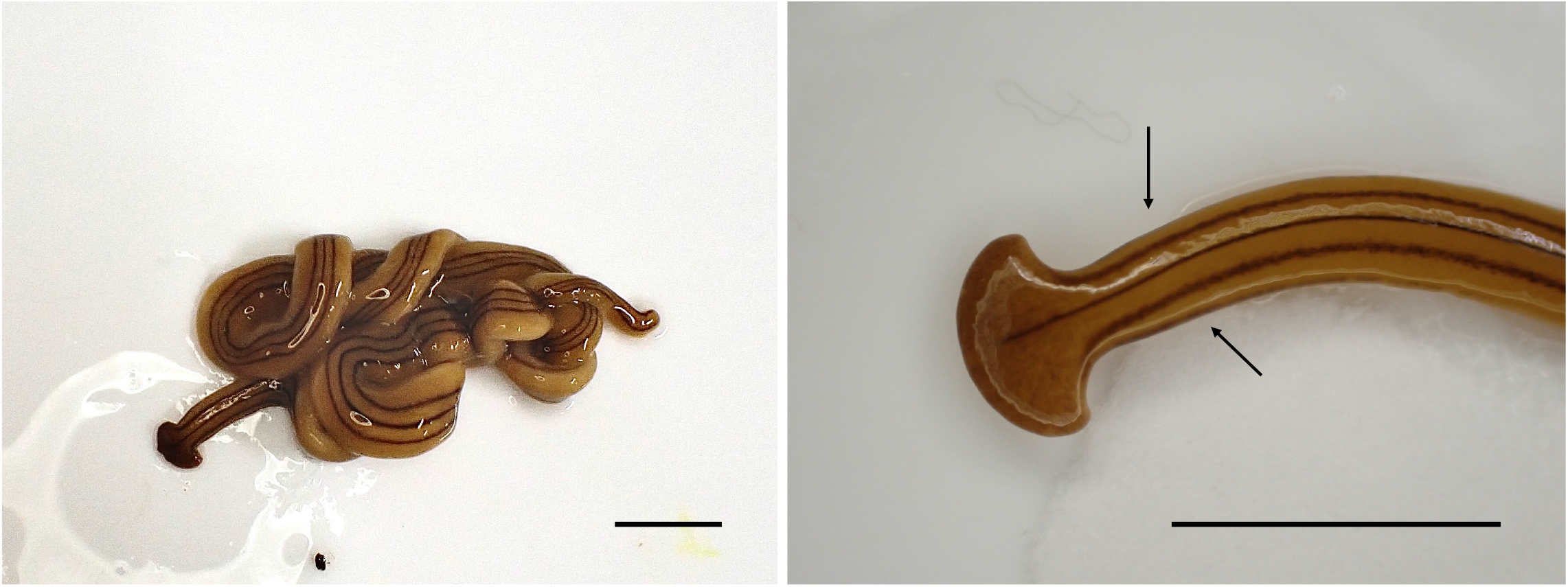
Photographs of the *Bipalium nobile* specimen analyzed in this study. (A) Whole body; (B) Head region. Three characteristic longitudinal stripes run along the entire length of the body. In the neck region, indicated by two arrows, two additional stripes appear along the dorsoventral boundaries, bringing the total number of stripes to five. Scale bars = 10 mm. The photographs were taken by the authors.

The mitogenome was assembled from the sequencing reads using NOVOPlasty v4.3.5 (Dierckxsens et al 2017) with the cox1 gene of *B. nobile* (GenBank: MG436936) as the seed. Gene annotation was conducted using MITOS2 WebServer (Al Arab et al. 2017; Donath et al. 2019) with the following options: Genetic code “Echinoderm, Flatworm (9)”, and Reference data “RefSeq63 Metazoa.” Inaccurate gene boundaries of rrnS, rrnL, and ND4 were corrected through visual observation. The genome map was drawn using OGDRAW (Greiner et al. 2019). Sequencing reads were mapped to the assembled mitogenome using BWA-MEM v0.7.17-r1188 (Li et al. 2013) and sorted using SAMtools v1.21 (Li et al. 2009). The sequencing depth across the entire mitogenome was calculated and visualized using the Python script deposited previously (Ni et al. 2023) (Figure S1).

Phylogenetic analysis was conducted based on amino acid sequences of all proteins encoded by the mitochondrial genomes of *B. nobile*, 17 other species belonging to the family Geoplanidae, and two species of the family Dugesiidae as outgroups. The sequences of species other than *B. nobile* were obtained from NCBI. Orthologous groups of 12 mitochondrial proteins across the 20 species were inferred using OrthoFinder v2.5.4 (Emms and Kelly 2019) and were subsequently used to construct the phylogenetic tree. The 12 protein sequences were aligned using MAFFT v7.525 (Katoh and Standley 2013), and non-aligned residual sequences were trimmed using trimAl v1.5.rev0 (Capella-Gutiérrez et al 2009). A maximum-likelihood (ML) tree was reconstructed based on the trimmed sequences using IQ-TREE v2.4.0 (Minh et al. 2020), employing 1000 bootstrap pseudoreplicates. The most appropriate amino acid substitution models for the 12 proteins were automatically selected by IQ-TREE. The chosen models are as follows: mtZOA+F+G4 for atp6, cob, cox3, ND2, ND3, ND6 and ND4L; mtZOA+I+R3 for cox1; mtZOA+F+I+G4 for cox2; mtZOA+F+I+R3 for ND1; and mtInv+F+I+G4 for ND4 and ND5.

## 3. Results

The mitochondrial genome of *B. nobile* sequenced in this study was 16,018 bp in length and contained 12 protein-coding genes, 22 transfer RNA (tRNA) genes, and 2 ribosomal RNA (rRNA) genes (Figure 2). The composition and order of genes were consistent with those observed in the closely related species *B. kewense* (Gastineau et al. 2019) and *D. multilineatum* (Justine et al. 2022), except for the position of the tRNA-Glu gene (Figure S2). In *B. kewense* and *D. multilineatum*, the tRNA-Glu is located between the tRNA-Lys and the tRNA-Ile, whereas in *B. nobile*, it has been relocated to a position between NAD5 and tRNA-Ser2 (Figure S2). There are three pairs of overlapping genes: tRNA-Asp-tRNA-Arg, tRNA-Leu1-tRNA-Tyr, and 16S rRNA-tRNA-Leu2 (Table 1). Due to the repetitive sequence in the control region, the genome could not be reconstructed as a circular sequence. To validate the mitochondrial genome sequence of *B. nobile* obtained in this study and to assess its consistency with previously proposed phylogenetic relationships, we inferred a phylogenetic tree using the maximum likelihood method based on amino acid sequences encoded by all mitochondrial protein-coding genes from *B. nobile* and other species in the family Geoplanidae for which mitochondrial genome sequences are available (Figure 3). The resulting tree placed *B. nobile* within a clade composed of members of the subfamily Bipaliinae. Within this clade, *B. nobile, B. kewense*, and *D. multilineatum* formed a monophyletic group.

**Table 1.** Gene content and composition of mitochondrial genome of *Bipalium nobile*.

| Feature | Type | Start | End | Length(bp) | Direction | Start codon | Stop codon |
| --- | --- | --- | --- | --- | --- | --- | --- |
| Control region | misc_feature | 15498/1 | 16018/227 | 748 | None | — | — |
| trnT | tRNA | 228 | 293 | 66 | Forward | — | — |
| trnN | tRNA | 308 | 385 | 78 | Forward | — | — |
| cob | gene | 391 | 1506 | 1116 | Forward | ATG | TAA |
| ND4L | gene | 1528 | 1815 | 288 | Forward | ATG | TAA |
| ND4 | gene | 1886 | 3334 | 1449 | Forward | ATG | TAA |
| trnF | tRNA | 3390 | 3459 | 70 | Forward | — | — |
| cox1 | gene | 3487 | 5037 | 1551 | Forward | ATG | TAG |
| trnE | tRNA | 5055 | 5126 | 72 | Forward | — | — |
| ND6 | gene | 5162 | 5650 | 489 | Forward | ATG | TAA |
| ND5 | gene | 5732 | 7312 | 1581 | Forward | ATT | TAA |
| trnQ | tRNA | 7584 | 7649 | 66 | Forward | — | — |
| trnS2 | tRNA | 7676 | 7744 | 69 | Forward | — | — |
| trnD | tRNA | 7753 | 7826 | 74 | Forward | — | — |
| trnR | tRNA | 7825 | 7890 | 66 | Forward | — | — |
| cox3 | gene | 7901 | 8686 | 786 | Forward | ATG | TAG |
| trnI | tRNA | 8697 | 8762 | 66 | Forward | — | — |
| trnK | tRNA | 8836 | 8904 | 69 | Forward | — | — |
| atp6 | gene | 8929 | 9597 | 669 | Forward | ATG | TAA |
| trnV | tRNA | 9604 | 9676 | 73 | Forward | — | — |
| ND1 | gene | 9720 | 10616 | 897 | Forward | ATT | TAG |
| trnW | tRNA | 10623 | 10699 | 77 | Forward | — | — |
| cox2 | gene | 10701 | 11369 | 669 | Forward | ATG | TAA |
| trnP | tRNA | 11376 | 11446 | 71 | Forward | — | — |
| ND3 | gene | 11477 | 11830 | 354 | Forward | GTG | TAA |
| trnA | tRNA | 11855 | 11923 | 69 | Forward | — | — |
| ND2 | gene | 11938 | 12927 | 990 | Forward | ATT | TAA |
| trnM | tRNA | 13103 | 13175 | 73 | Forward | — | — |
| trnH | tRNA | 13199 | 13271 | 73 | Forward | — | — |
| trnC | tRNA | 13293 | 13355 | 63 | Forward | — | — |
| rrnS | rRNA | 13356 | 14110 | 755 | Forward | — | — |
| trnL1 | tRNA | 14110 | 14188 | 79 | Forward | — | — |
| trnY | tRNA | 14179 | 14244 | 66 | Forward | — | — |
| trnG | tRNA | 14268 | 14326 | 59 | Forward | — | — |
| trnS1 | tRNA | 14356 | 14419 | 64 | Forward | — | — |
| rrnL | rRNA | 14420 | 15486 | 1067 | Forward | — | — |
| trnL2 | tRNA | 15430 | 15497 | 68 | Forward | — | — |

**Figure 2.**
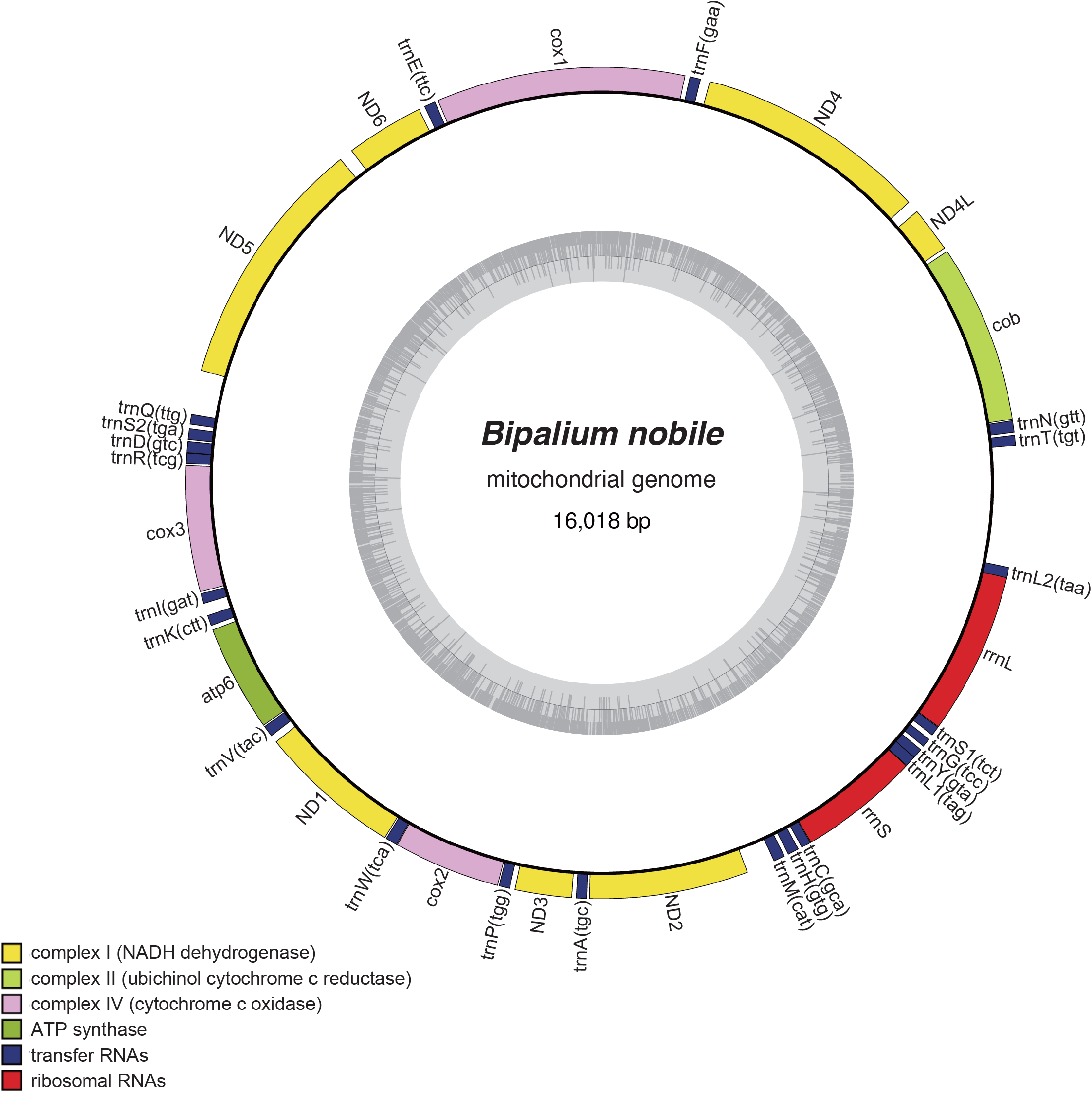
Mitochondrial genome map of *Bipalium nobile*. Although the sequence obtained in this study was not circular, it is presented here as a circular map. The gap in the sequence is located between tRNA-Thr and tRNA-Leu2.

**Figure 3.**
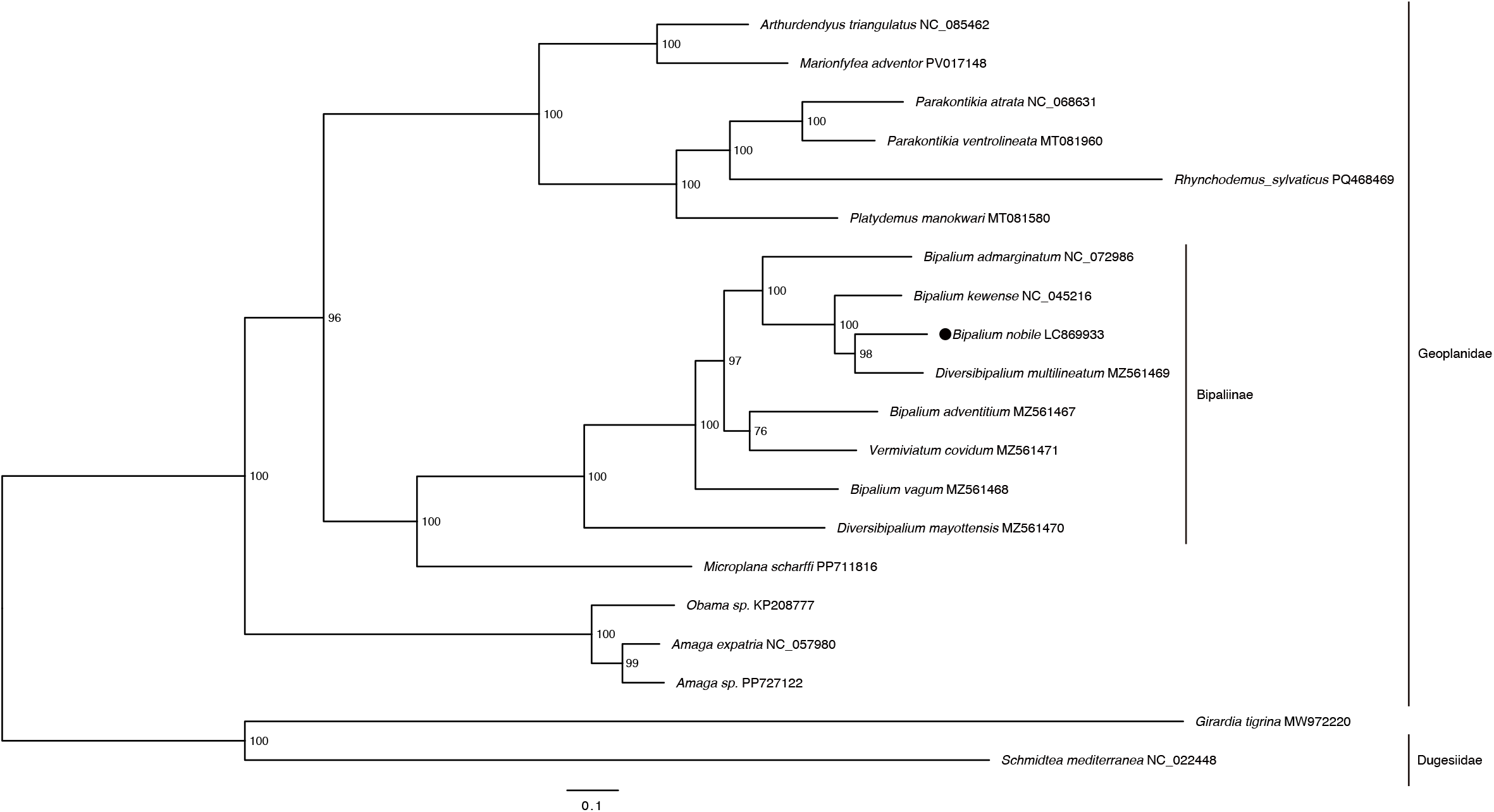
Phylogenetic tree of *Bipalium nobile* and other species in the family Geoplanidae inferred using the maximum likelihood method based on the amino acid sequences of all proteins encoded by mitochondrial genomes. Numbers at nodes indicate bootstrap values. The scale bar represents 0.1 substitutions per site. *B. nobile* is highlighted by a black dot. *Girardia tigrina* and *Schmidtea mediterranea* were included as outgroups. The references for the sequences used are as follows: *Amaga expatria* NC_057980 (Justine et al 2020); *Amaga sp*. PP727122 (unpublished); *Arthurdendyus triangulatus* NC_085462 (unpublished); *Bipalium admarginatum* NC_072986 (unpublished); *Bipalium adventitium* MZ561467 (Justine et al 2022); *Bipalium kewense* NC_045216 (Gastineau et al 2019); *Bipalium nobile* LC869933 (this study); *Bipalium vagum* MZ561468 (Justine et al 2022); *Diversibipalium mayottensis* MZ561470 (Justine et al 2022); *Diversibipalium multilineatum* MZ561469 (Justine et al 2022); *Girardia tigrina* MW972220 (unpublished); *Marionfyfea adventor* PV017148 (unpublished); *Microplana scharffi* PP711816 (unpublished); *Obama sp*. KP208777 (Solà et al 2015); *Parakontikia atrata* NC_068631 (Gastineau et al 2022); *Parakontikia ventrolineata* MT081960 (Gastineau and Justine 2020); *Platydemus manokwari* MT081580 (Gastineau et al 2020); *Rhynchodemus sylvaticus* PQ468469 (unpublished); *Schmidtea mediterranea* NC_022448 (unpublished); *Vermiviatum covidum* MZ561471 (Justine et al 2022).

## 4. Discussion and Conclusion

To date, three mitochondrial gene sequences of *B. nobile* have been registered in public databases, all corresponding to the cox1 gene: two derived from specimens collected in Japan (Morffe et al. 2020) and one from a specimen collected in Korea. When these sequences were compared with the cox1 gene sequence obtained in the present study, the approximately 750 bp sequences from the Japanese specimens were found to be identical to that of this study, whereas the sequence from the Korean specimen differed at 3 positions out of 387 base pairs (Figure S3). These results suggest the presence of genetic diversity between the Japanese and Korean populations of *B. nobile*. The mitochondrial genome sequence obtained in this study is expected to serve as a valuable resource for future research aimed at elucidating the genetic diversity of *B. nobile*.

Solà et al. (2023) reported a phylogenetic tree of the subfamily Bipaliinae based on the mitochondrial cox1 gene and the nuclear 28S and 18S rRNA genes. Although the gene sets and operational taxonomic units used in their analysis differ from those in the present study, both studies consistently recovered a monophyletic clade comprising *B. nobile, B. kewense*, and *D. multilineatum*. Solà et al. (2023) proposed that this monophyletic group should be recognized as a new genus. The findings of the present study support the close phylogenetic relationship among these three species and demonstrate the similarity in mitochondrial genome organization shared by them. Further sequencing of mitochondrial genomes from additional species within the subfamily Bipaliinae will allow for a more thorough assessment of the validity of this proposal.

## 5. Ethical approval

The material of this paper does not involve ethical conflicts. *B. nobile* is neither endangred on the CITES catalog nor collected from a natural reserve.

## 6. Author contributions

MO designed the experiment, collected the specimen, extracted genomic DNA, annotated the mitochondrial genome, and wrote the manuscript. ST annotated the mitochondrial genome, constructed phylogenetic tree, and wrote the manuscript. RM assembled mitochondrial genome. DH handled the procedures for specimen deposition and assembled mitochondrial genome. SW designed and conducted the experiment and wrote the manuscript.

## 9. Data availability statement

The mitochondrial genome sequence of *B. nobile* is openly available in GenBank of NCBI at https://www.ncbi.nlm.nih.gov, under the accession no. LC869933. The associated BioProject, SRA, and Bio-Sample numbers are PRJDB20669, DRX645385, and SAMD00902039, respectively.

## Legends for supplementary figures

**Figure S1.**
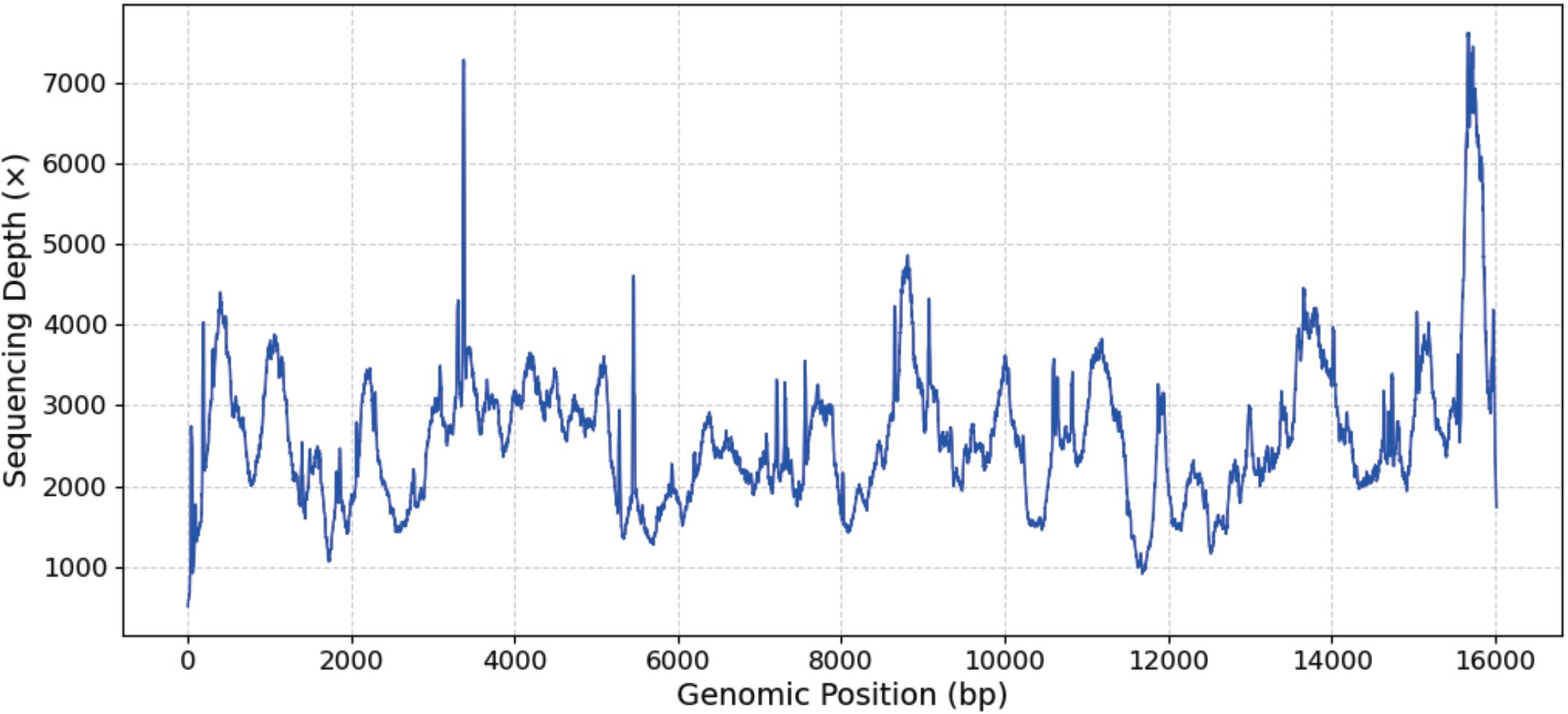
Read coverage depth map of the mitochondrial genome of *Bipalium nobile*. Average, maximal, and minimal depths were 2607.18, 7619, and 506, respectively.

**Figure S2.**
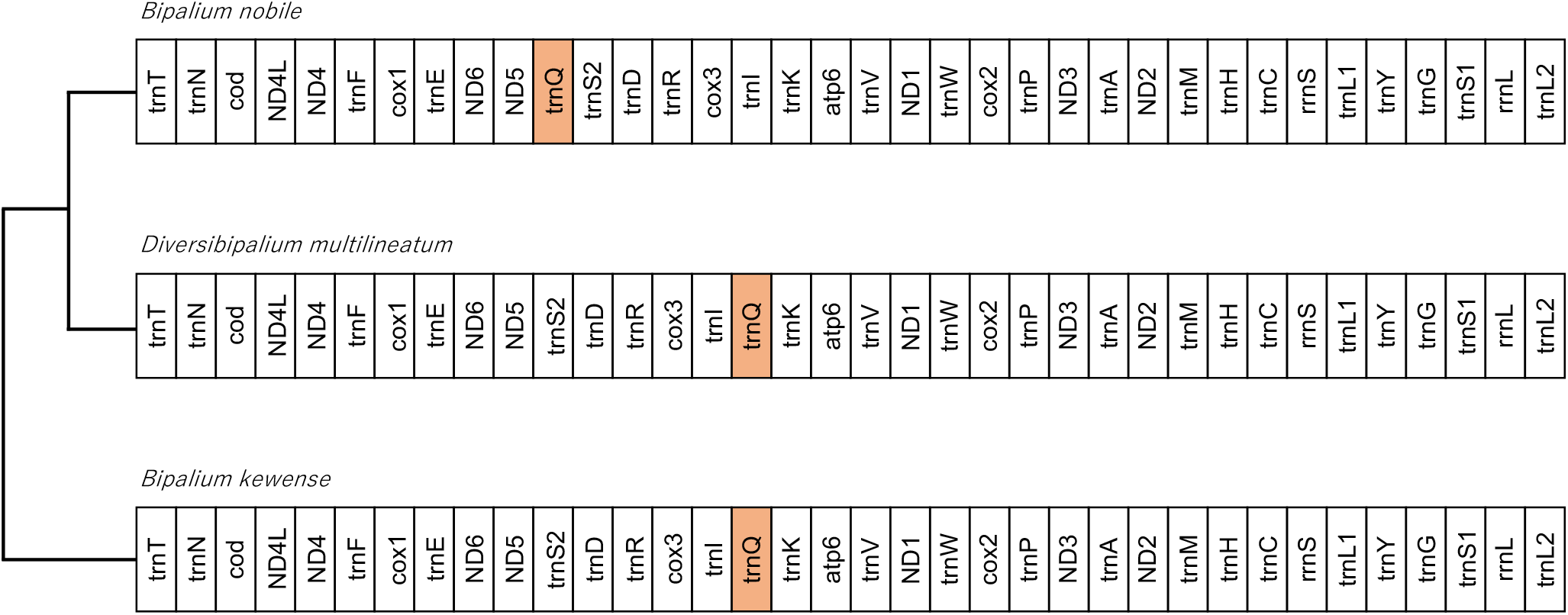
Comparison of mitochondrial gene order in *Bipalium nobile* and related species. The gene order in the mitochondrial genomes of *B. nobile, Bipalium kewense*, and *Diversibipalium multilineatum* is compared. Except for the position of the tRNA-Glu gene, shown in orange, the composition and order of genes are conserved.

**Figure S3.**
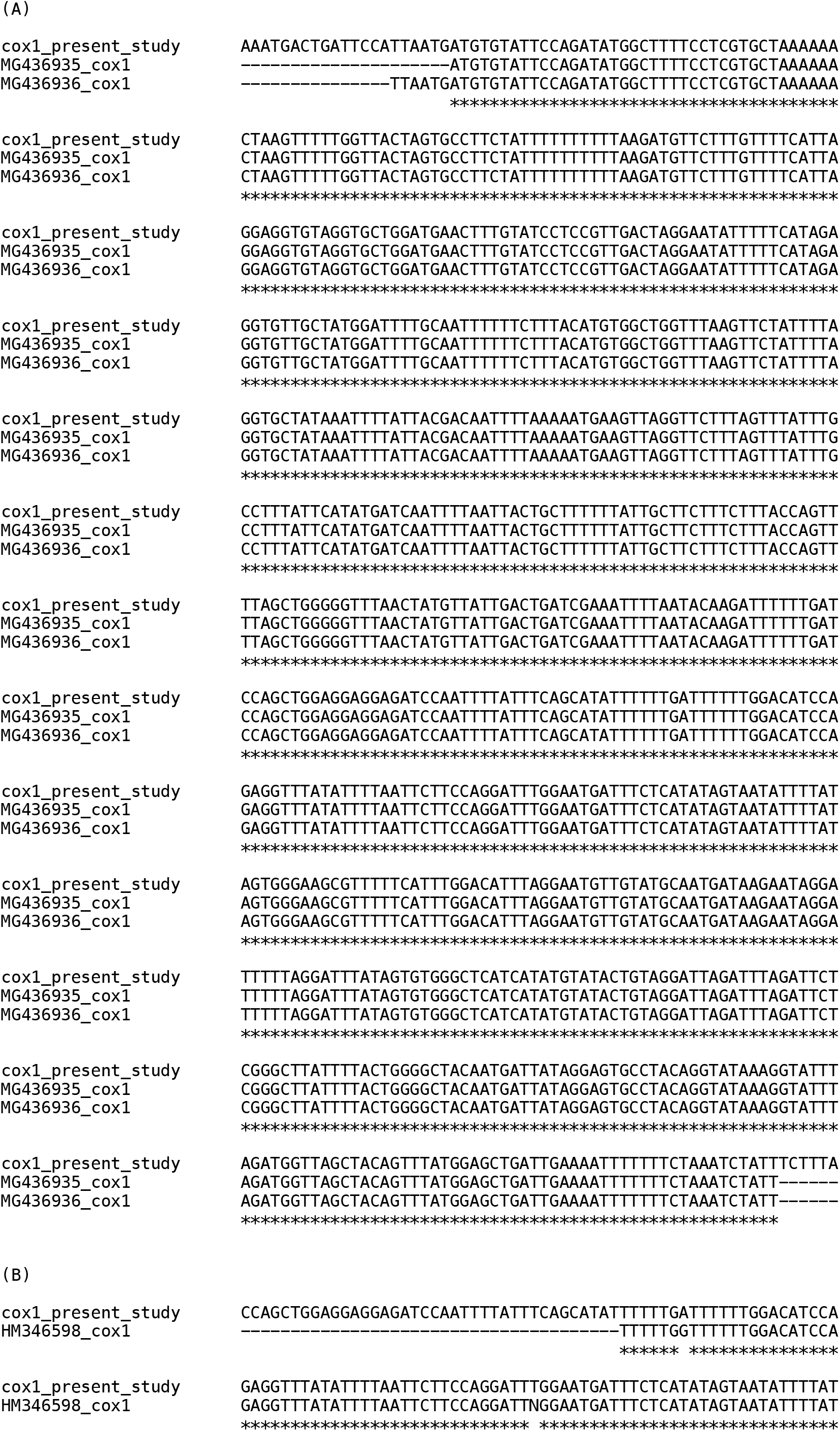
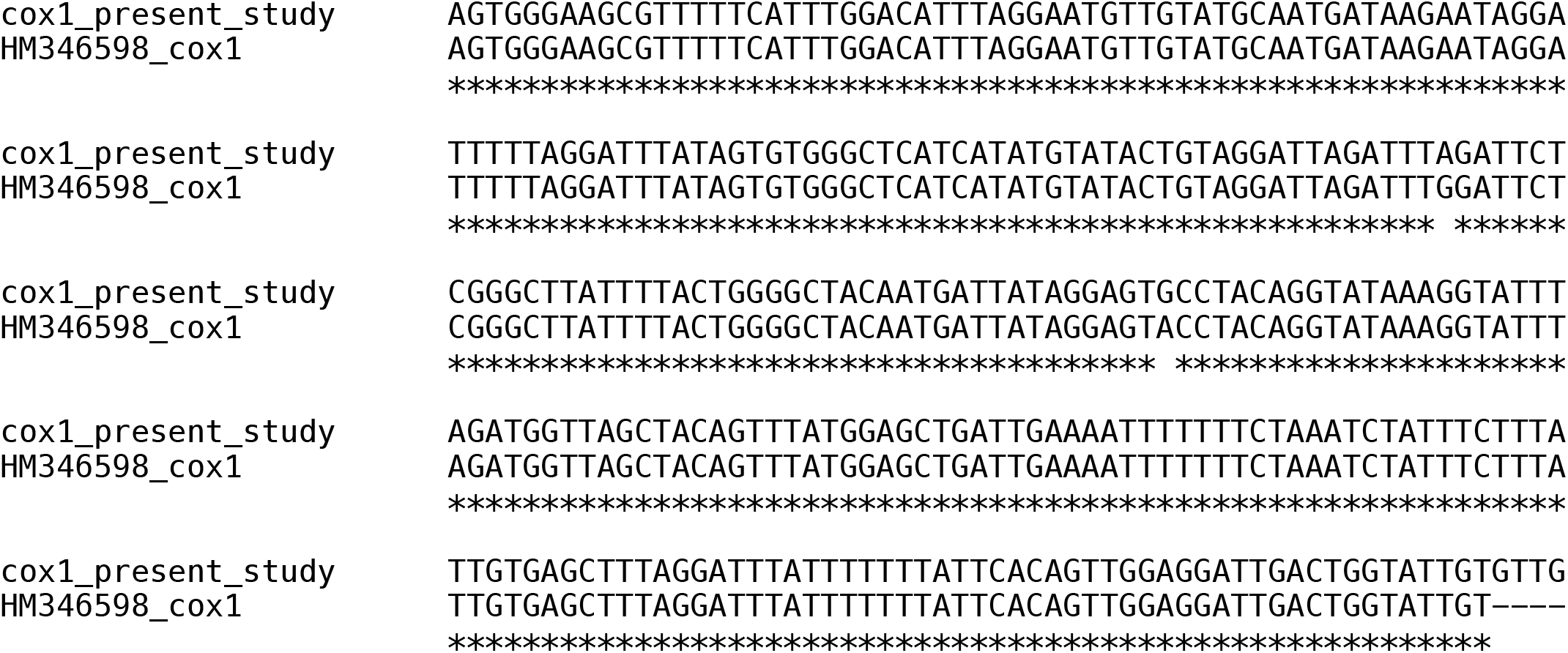
Comparison of nucleotide sequences in the mitochondrial cox1 gene region of *Bipalium nobile*. (A) Comparison between the sequence obtained in the present study and two previously registered sequences from Japanese specimens (MG436935 and MG436936; Morffe et al. 2020). (B) Comparison between the sequence obtained in the present study and one previously registered sequence from a Korean specimen (HM346598; unpublished). An asterisk indicates identical nucleotides.

